# Adaptive Immunity Governs Regional Aortic Remodeling in Hypertension via Perivascular Adipose Tissue Plasticity

**DOI:** 10.1101/2025.07.23.666477

**Authors:** Yujun Xu, J. Caleb Snider, Matthew R. Bersi

## Abstract

Hypertension drives heterogeneous aortic remodeling, but the mechanisms underlying regional disparities remain unclear. Here, we demonstrate that adaptive immunity orchestrates spatial differences in vascular dysfunction by modulating perivascular adipose tissue (PVAT) phenotype and immune-metabolic crosstalk. Using angiotensin II (AngII)-infused wild-type (WT) and *Rag1^−/−^*mice lacking T and B cells, we integrated biaxial mechanical testing, bulk transcriptomics, and PVAT analyses. In WT mice, AngII induced pronounced descending thoracic aorta (DTA) remodeling, marked by wall thickening, reduced circumferential stiffness and inflammatory gene upregulation (*Il6*, *Ccl2*). These changes were attenuated in *Rag1^−/−^* mice, implicating T cells in thoracic maladaptation. Conversely, the infrarenal abdominal aorta (IAA) exhibited hypertensive resilience in WT mice but unmasked PPARγ-associated metabolic reprogramming (*Pparg*, *Adipoq*) in *Rag1^−/−^* mice, suggesting T cells suppress protective abdominal adaptations. PVAT heterogeneity emerged as a key regulator wherein thoracic PVAT (T-PVAT) adopted a pro-inflammatory phenotype (CCL5, TIMP-1) in WT mice, exacerbating DTA damage, while *Rag1^−/−^* mice showed thermogenic plasticity (*Ucp1* upregulation) in abdominal PVAT (A-PVAT). T cell reconstitution restored maladaptive remodeling in *Rag1^−/−^* mice, confirming adaptive immunity’s dual role in promoting thoracic injury and restraining metabolic resilience. This work identifies PVAT as an immune-metabolic switch governing regional susceptibility to vascular remodeling, offering spatially resolved strategies to preserve aortic compliance in hypertensive disease.

## Introduction

Hypertension, a leading cause of cardiovascular morbidity and mortality, affects ∼48.1% of U.S. adults and contributes to ∼685,000 deaths annually^1^. Arterial stiffening, a hallmark of vascular disease, impairs aortic capacity to buffer pulsatile pressure, thus contributing to hemodynamic dysfunction and promoting end-organ damage^2^. This pathological remodeling is multi-factorial, arising from dysregulated interactions between hemodynamic stress, extracellular matrix (ECM) degradation, collagen deposition, and immune activation^3,4^.

The aorta exhibits regional heterogeneity in its susceptibility and response to chronic hypertension. Biomechanical studies have revealed pronounced stiffening and ECM remodeling in the descending thoracic aorta (DTA) compared to the infrarenal abdominal aorta (IAA)^5^. These disparities correlate with increased adventitial collagen accumulation and inflammatory cell infiltration in the DTA^5–7^, yet the mechanisms driving spatial differences in immuno-mechanobiological responses remain unresolved.

Adaptive immunity, particularly T cell activity, is a critical mediator of hypertensive vascular injury. T cells are recruited to the perivascular space, infiltrate the adventitia, and release pro-inflammatory cytokines (e.g., IL-6, IL-17A) that drive macrophage recruitment, collagen deposition, and endothelial dysfunction^7,8^. While immune-mediated thoracic aortic remodeling is well-documented^9^, the IAA’s resilience and its potential metabolic adaptations to hypertension remain poorly characterized.

Perivascular adipose tissue (PVAT) surrounds blood vessels and acts as an endocrine organ to regulate vascular tone and inflammation through adipokine secretion and immune cell crosstalk^10–15^. PVAT also exhibits anatomical phenotypic diversity. Specifically, thoracic PVAT (T-PVAT) displays brown adipose-like properties, including high *Ucp1* expression and thermogenic capacity; in contrast, abdominal PVAT (A-PVAT) resembles white adipose tissue, exhibiting high lipid content and pro-inflammatory characteristics^16–18^. This phenotypic gradient suggests PVAT heterogeneity may be associated with regional vascular responses to hypertension. For instance, T-PVAT inflammation exacerbates endothelial dysfunction^12^, while A-PVAT derived adiponectin enhances vasodilation^10,19^. Furthermore, T cells recruited to PVAT depots can dynamically regulate vascular stiffening; the pro-inflammatory Th1/Th17 subsets drive perivascular fibrosis and Foxp3⁺ regulatory T cells secrete IL-10 to preserve thermogenic function and attenuate vessel stiffening^10,20,21^. T cell activity can also directly modulate PVAT phenotype. Namely, PVAT-derived chemokines (e.g., CCL5) potently recruit T cells to the perivascular space^12^ and IFNγ-secreting Th1 cells suppress thermogenic gene expression (*Ucp1, Pgc1α*)^22^, thus potentiating a synergistic feedback loop.

We therefore hypothesized that adaptive immunity may differentially regulate PVAT phenotype, structure, and function across different aortic regions, ultimately driving spatial disparities in vascular remodeling. Using angiotensin II (AngII)-infusion model of induced hypertension in wild-type (WT) and *Rag1*^-/-^ mice lacking mature lymphocytes (e.g., T and B cells), we investigated this hypothesis through integrated biaxial mechanical testing, transcriptomic profiling, and PVAT analyses. We further hypothesized that T cells mediate maladaptive thoracic remodeling through excess inflammation and ECM degradation and modulate metabolic pathways to promote adaptation in the abdominal aorta. Thus, T-PVAT adopts a pro-inflammatory phenotype during hypertension that exacerbates thoracic damage, whereas A-PVAT exhibits enhanced metabolic plasticity linked to preserved vascular function.

This work advances prior mechanocentric models^5^ by identifying PVAT heterogeneity as an immune-metabolic switch governing regional susceptibility to hypertensive vascular remodeling. The distinct molecular signatures of T-PVAT versus A-PVAT create microenvironments that critically shape local vascular responses to hypertension. These insights provide a foundation for PVAT-targeted therapies to preserve aortic compliance and mitigate hypertensive complications.

## Methods

Detailed information regarding all materials and methods used in the current study, including experimental data collection and mathematical modeling, are available in the Supplemental Material. A brief description of the statistical analysis approach is described below.

### Statistical analysis

Statistical analyses were performed using GraphPad Prism (v10.0; GraphPad Software, San Diego, CA). Data are presented as mean ± standard error of the mean (SEM), with individual data points representing biological replicates. Sample sizes for each experiment are detailed in figure legends. For comparisons between two groups, unpaired two-tailed Student’s *t*-tests were applied to datasets with equal variances. For multi-group comparisons involving a single variable, unpaired two-tailed Student’s *t* test was employed; ordinary two-way ANOVA followed by Tukey’s multiple comparisons test was utilized for comparisons with two factors. Statistical significance was defined as *p* ≤ 0.05 with group-specific differences indicated in individual figures where appropriate.

## Results

### Lymphocyte-Mediated Regional Biomechanical Remodeling in AngII-Treated Aortas

To investigate the spatial interplay between mature lymphocytes and biomechanical remodeling in hypertensive aortas, we compared thoracic (DTA) and abdominal (IAA) aortic responses from WT and lymphocyte-deficient *Rag1*^⁻/⁻^ mice following 14 days of AngII infusion. In WT mice, AngII induced significant wall thickening in the DTA under both unloaded and loaded (120 mmHg systolic pressure) conditions (**Fig. 1a, left**; +14.50% unloaded and +20.26% loaded; *p* < 0.0001), whereas the IAA showed no detectable changes (**Fig. 1a, middle-left**; *p* > 0.05). This regional disparity persisted but was attenuated in *Rag1*^⁻/⁻^ mice: AngII elicited only marginal wall thickening in *Rag1*^⁻/⁻^-DTA (**Fig. 1a, middle-right**; +9.57% unloaded and +18.50% loaded; unloaded: *p* = 0.059; loaded: *p* = 0.053), while *Rag1*^⁻/⁻^-IAA remained unaffected (**Fig. 1a, right**; *p* > 0.05). These findings implicate adaptive immunity as a contributor to region-specific aortic remodeling, with the DTA exhibiting greater immune-mediated structural adaptation.

**Fig. 1.**
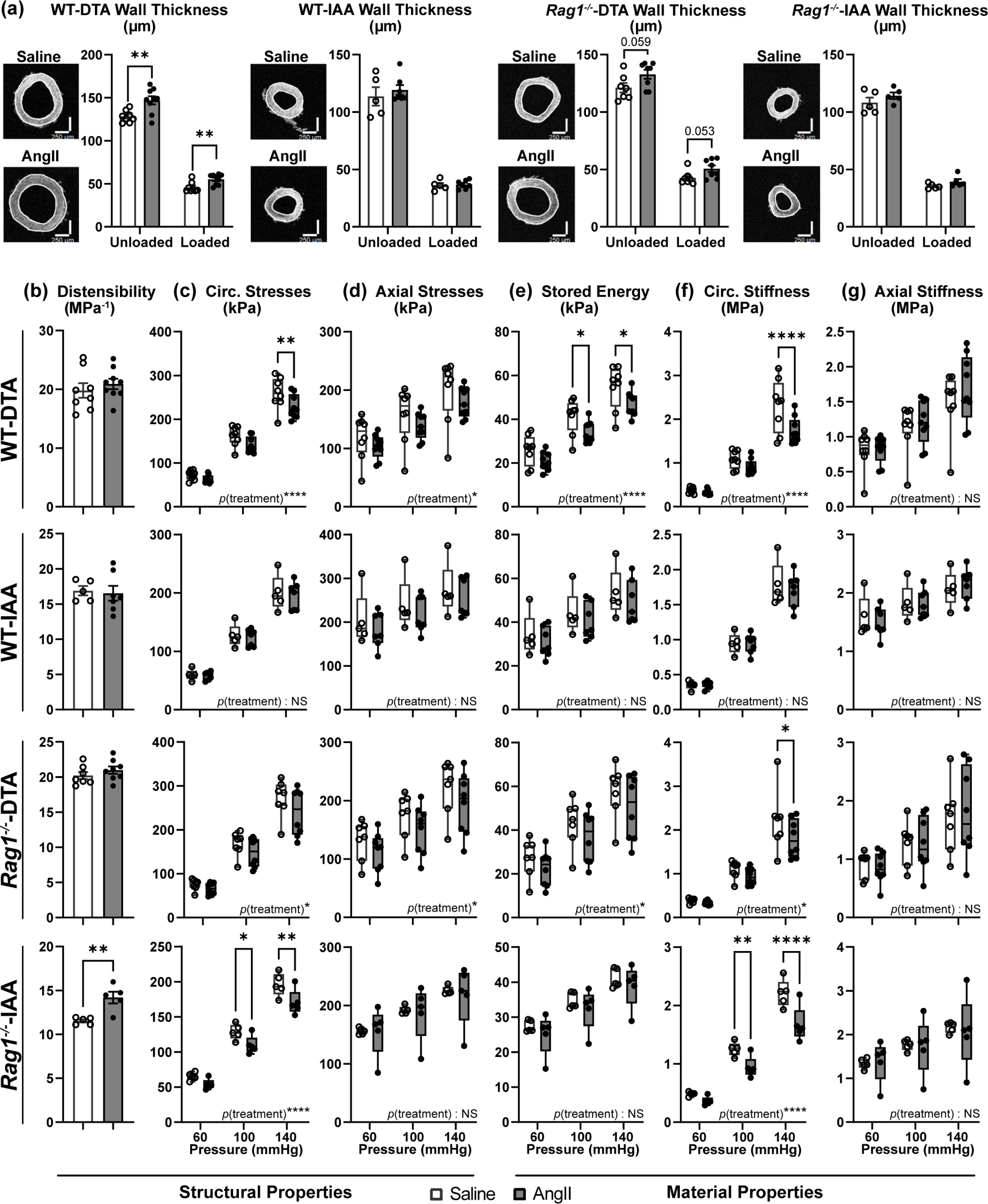
Immune-mediated regional biomechanical remodeling in hypertensive aortas. **(a)** unloaded and loaded wall thickness (µm), **(b)** distensibility (MPa⁻¹), **(c,d)** circumferential and axial stresses (kPa), **(e)** stored energy (kPa), and **(f,g)** circumferential and axial stiffness (MPa) in DTA and IAA of WT and *Rag1^⁻/⁻^* mice following 14 days of Saline or AngII infusion. *p_blood pressure* ≤ 0.0001 for all groups (b-g). See **Fig. S1** for sample sizes, calculated properties at systolic pressure (120 mmHg) and best-fit parameters. Scale bar = 250µm. All data shown as mean ± SEM. Statistical significance from either (a) unpaired two-tailed Student’s *t*-tests or (b-g) two-way ANOVA with post hoc testing for multiple comparisons denoted by NS (*p*>0.5), \**p* ≤ 0.05, \*\**p* ≤ 0.01, \*\*\**p* ≤ 0.001, or \*\*\*\**p* ≤ 0.0001.

To assess regional responses to pressurization, we calculated distensibility – a metric inversely related to structural stiffness – using a four-fiber family constitutive model. At an estimated systolic pressure (120 mmHg), AngII significantly increased distensibility in *Rag1*^⁻/⁻^-IAA (**Fig. 1b**, **S1a–d**; *p* < 0.001), whereas WT-DTA, WT-IAA, and *Rag1*^⁻/⁻^-DTA retained near-basal distensibility (**Fig. 1b**; *p* > 0.05). Elevated distensibility in *Rag1*^⁻/⁻^-IAA suggests reduced structural stiffness, likely reflecting compensatory AngII-induced remodeling in the absence of T cells. Consistent with these findings, circumferential stresses, calculated from the pressure and corresponding geometry (see Supplementary Methods), dropped markedly in *Rag1*^⁻/⁻^-IAA at 100 and 140 mmHg due to AngII-treatment (**Fig. 1c**; *p_treatment* < 0.0001, two-way ANOVA). This decrease in stress, despite unchanged wall thickness, implies altered load-bearing capacity. In contrast, WT-DTA exhibited reduced circumferential stresses at 140 mmHg under AngII (**Fig. 1c**; *p_treatment* < 0.0001), correlating with its thickened wall; WT-IAA showed no significant changes. Axial stresses – a measure of structural integrity in the longitudinal direction – remained unaffected across groups at individual pressures (*p_treatment* > 0.05), although two-way ANOVA revealed significant treatment effects in WT- and *Rag1*^⁻/⁻^-DTA (**Fig. 1d**; *p_treatment* < 0.05). These data underscore distinct directional and regional adaptations in response to hypertensive stimulus.

Estimation of material properties and model-derived mechanical metrics further highlighted regional specificity. Stored energy – a marker of mechanical energy retention and a hallmark of vascular function – declined significantly in WT-DTA near systolic pressures following AngII infusion (**Fig. 1e**; *p_treatment* < 0.05), indicating impaired mechanical resilience. Global treatment effects were detected in *Rag1*^⁻/⁻^-DTA (*p_treatment* < 0.05, two-way ANOVA), though pairwise comparisons showed no significance. Circumferential material stiffness, a measure of resistance to change in aortic diameter, was significantly reduced in AngII-treated WT-DTA (140 mmHg; *p_treatment* < 0.0001), *Rag1*^⁻/⁻^-DTA (140 mmHg; *p* < 0.05), and *Rag1*^⁻/⁻^-IAA (100–140 mmHg; *p_treatment* < 0.001), but not in WT-IAA (**Fig. 1f**; *p_treatment* > 0.05). Reduced stiffness in these regions suggests potential ECM degradation or collagen disorganization as a consequence of AngII infusion. In contrast, axial material stiffness remained unchanged in all groups (*p_treatment* > 0.05). These findings elucidate direction-specific extracellular matrix adaptations throughout the aorta in response to AngII. Note that applied blood pressure exhibited universal effects on biomechanical metrics (*p*_*blood pressure* < 0.0001) in all groups, confirming the intrinsic pressure-responsive behavior of the aorta.

Together, these results demonstrate that AngII-induced aortic remodeling exhibits region-specific effects, with the thoracic aorta demonstrating immune-mediated susceptibility and the abdominal aorta maintaining structural integrity and mechanical resilience. In *Rag1*^⁻/⁻^ mice, the IAA showed compensatory increases in distensibility, suggesting an adaptive ECM remodeling in the absence of T cells. These findings indicate adaptive immunity is a critical regulator of spatially heterogeneous vascular remodeling during hypertension.

### Regional Transcriptomic Profiling Reveals Immune-Mediated Mechanisms of Aortic Remodeling during Hypertension

Following biomechanical analysis, we conducted comprehensive transcriptomic profiling to explore the molecular mechanisms underlying region-specific aortic remodeling in WT and *Rag1^-/-^* mice. Expression of key genes revealed distinct signatures across aortic regions and genotypes under AngII treatment. We categorized differentially expressed genes into four functional groups: immune-associated, vascular remodeling, adipogenic-associated, and bridge genes that link immune and metabolic pathways. In WT-DTA, AngII induced a robust inflammatory signature, marked by pronounced upregulation of immune-associated genes (*Il6*, *Il17ra*, *Cd3e*, *Ccl2,* etc.) and chemokines (*Ccl6*), alongside downregulation of the regulatory T cell marker *Foxp3* (**Fig. 2a**). This transcriptional profile coincides with prominent wall thickening and reduced circumferential stiffness in WT-DTA, as observed in the biomechanical analysis. Concurrently, vascular remodeling genes (*Mmp2*, *Timp1,* etc.) were upregulated in WT-DTA, while structural markers (*Acta2*, *Myh11,* etc.) declined, reflecting active ECM turnover. In contrast, WT-IAA exhibited minimal transcriptional changes after AngII treatment, consistent with its preserved biomechanical integrity. Notably, genes associated with adipogenesis remained largely unchanged in both aortic regions of WT mice after AngII treatment.

**Fig. 2.**
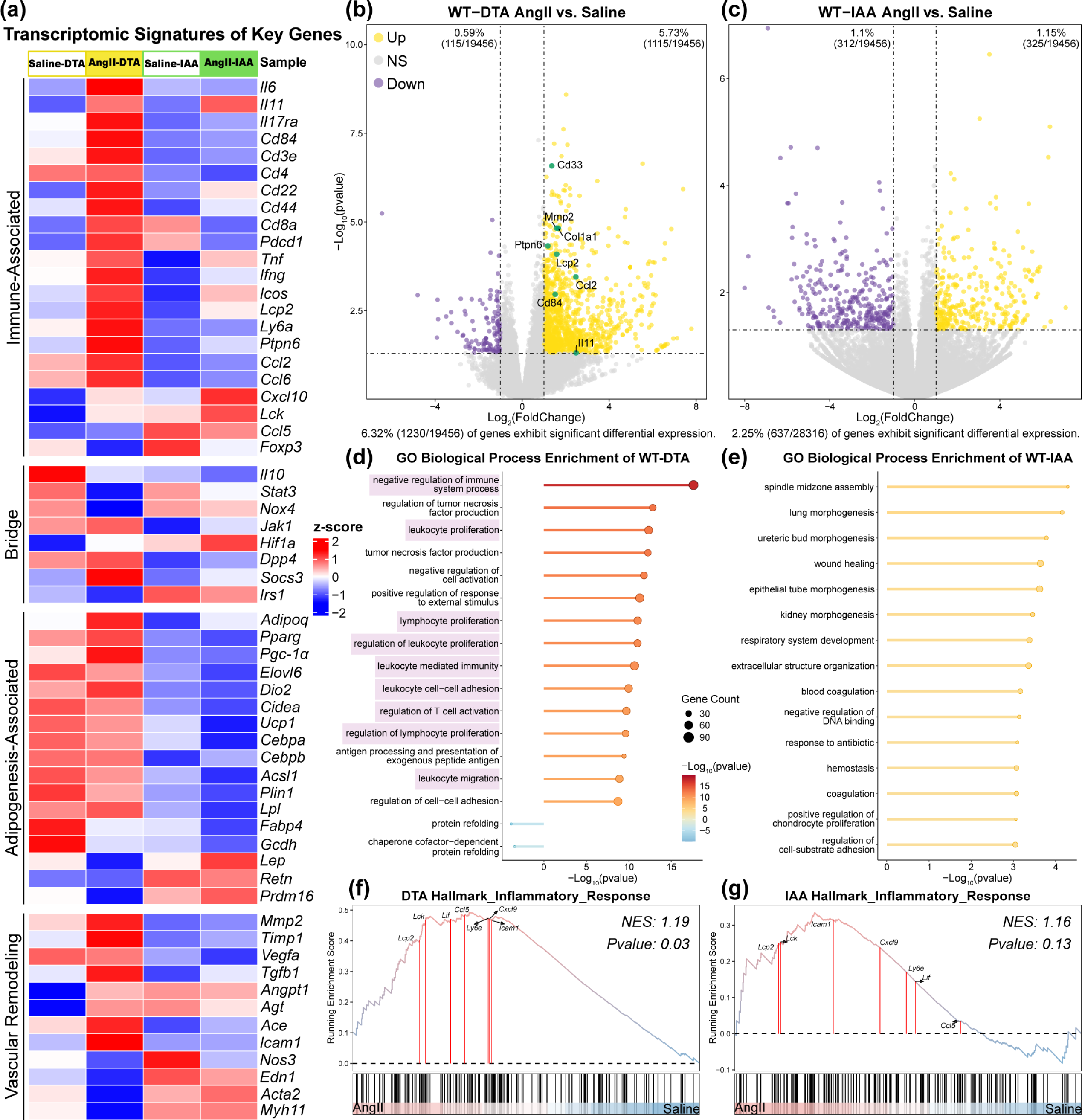
Region-specific immune-driven transcriptional changes in WT AngII-treated aortas. **(a)** Expression heatmap of key genes across immune regulation, adipogenesis, vascular remodeling and structural integrity in WT-DTA and WT-IAA (avg_cpm values) in response to Saline or AngII treatment. **(b,c)** Volcano plots of differential gene expression in WT-DTA (**b**; 1230 DEGs) and WT-IAA (**c**; 637 DEGs). Upregulated genes are highlighted in yellow; downregulated genes are highlighted in purple. Dashed line: *p* = 0.05, log₂(FoldChange) = ±1. **(d,e)** Top enriched GO biological process terms from WT-DTA and WT-IAA. Immune-related pathways are highlighted in purple. **(f,g)** GSEA enrichment plots of inflammatory response Hallmark pathway in WT-DTA and WT-IAA. n = 5 mice/group. See **Fig. S2** for top 15 Hallmark pathways in each group comparison.

Regional specificity in transcriptional responses was evident from differential gene expression patterns (**Fig. 2b,c**). In WT-DTA, 6.32% of genes were differentially expressed (DEG) after AngII infusion, with 5.73% of genes being markedly <u>upregulated,</u> including immune-related transcripts such as *Il6*, *Il17ra*, and *Ccl2* (**Fig. 2b**). In contrast, WT-IAA exhibited a subdued transcriptional response (2.25% DEG) following AngII, with balanced up- and downregulation (**Fig. 2c**). This threefold disparity highlights the DTA’s heightened transcriptional sensitivity and biological response to hypertensive stimulus, aligning with its biomechanical vulnerability.

Gene Ontology (GO) enrichment analysis confirmed region-specific transcriptional programs. In WT-DTA, 9 of the top 15 enriched biological processes were immune-related, including *immune regulation*, *leukocyte proliferation*, and *lymphocyte proliferation*, among others (purple; **Fig. 2d**). This immune dominance contrasted sharply with WT-IAA, where enriched pathways centered on tissue development and extracellular matrix organization, such as *wound healing* and *epithelial morphogenesis*, which may suggest an adaptive transcriptional signature in response to AngII (**Fig. 2e**). Gene Set Enrichment Analysis (GSEA) reinforced these regional disparities. The inflammatory response Hallmark pathway was significantly enriched in WT-DTA (NES: 1.19, *p* = 0.03), driven by T cell signaling components (*Lcp2*, *Lck*), chemokines (*Ccl5*, *Cxcl9*), and adhesion molecules (*Icam1*) (**Fig. 2f**); WT-IAA showed no significant enrichment (NES: 1.16, *p* = 0.13), underscoring its limited inflammatory response after AngII (**Fig. 2g**).

To further examine the role of adaptive immunity in AngII-mediated aortic remodeling, we performed parallel transcriptomic analysis in *Rag1^⁻/⁻^* mice. The absence of mature lymphocytes significantly altered expression of immune-associated genes (**Fig. 3a**). Regional transcriptional differences persisted but were attenuated compared to WT. Strikingly, adipogenesis-associated genes showed a trend toward upregulation in both *Rag1^⁻/⁻^*-DTA and *Rag1^⁻/⁻^*-IAA under AngII treatment, which was not observed in WT mice.

**Fig. 3.**
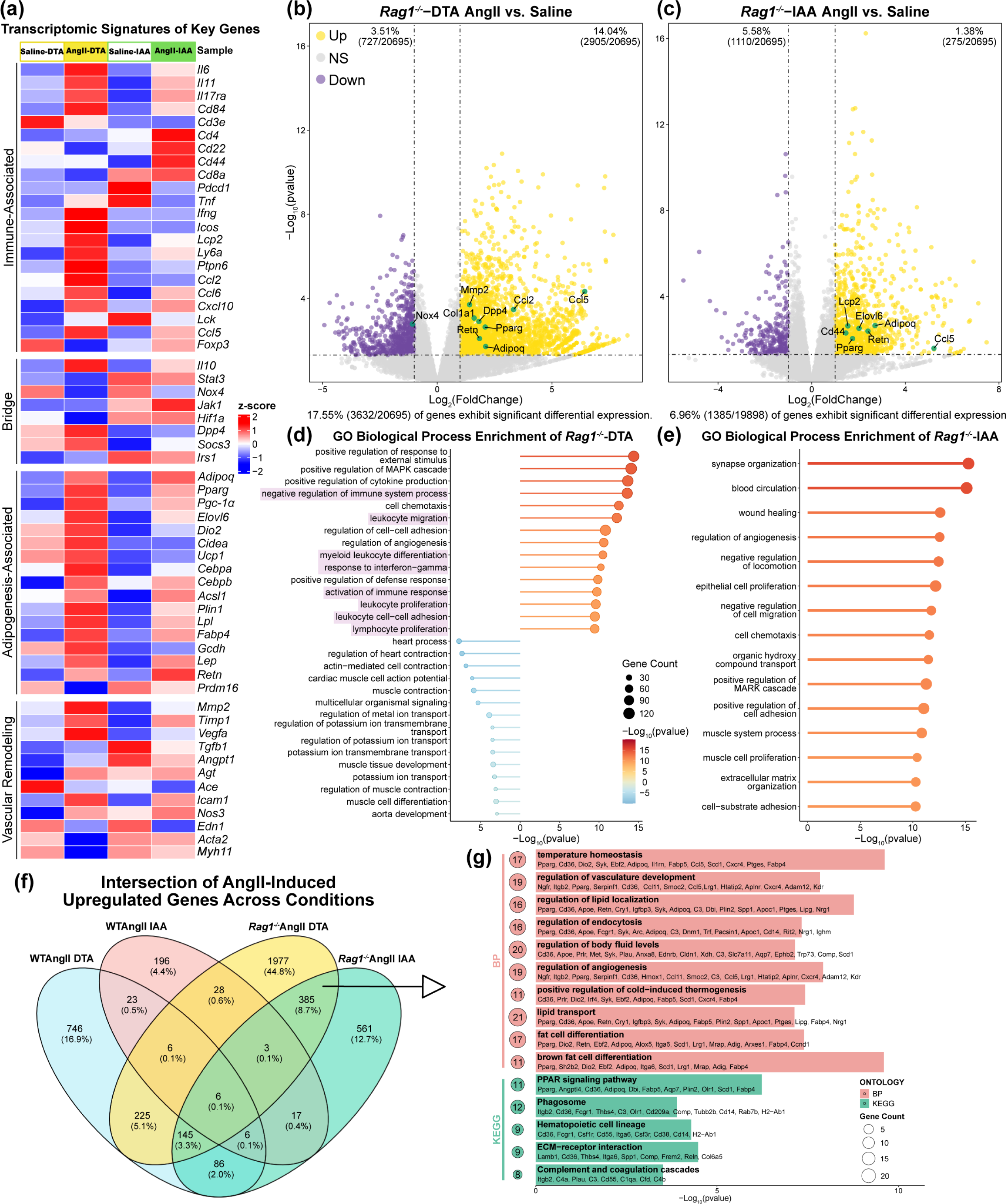
Region-specific compensatory transcriptional programs in lymphocyte-deficient AngII-treated aortas. **(a)** Expression heatmap of key genes across immune regulation, adipogenesis, vascular remodeling and structural integrity in *Rag1^-/-^*-DTA and -IAA (avg_cpm values) in response to Saline or AngII treatment. **(b,c)** Volcano plots of differential gene expression in *Rag1^-/-^*-DTA (**b**; 3632 DEGs) and -IAA (**c**; 1385 DEGs). Upregulated genes are highlighted in yellow; downregulated genes are highlighted in purple. Dashed line: *p* = 0.05, log₂(FoldChange) = ±1. **(d,e)** Top enriched GO biological process terms in *Rag1^-/-^*-DTA and -IAA. Immune-related pathways are highlighted in purple. **(f)** Venn diagram of upregulated DEGs across aortic regions and genotypes following AngII treatment. **(g)** Top enriched GO biological processes (pink) and KEGG pathways (green) for shared upregulated gene set between AngII-treated *Rag1^-/-^* regions (*Rag1^-/-^* AngII-DTA and *Rag1^-/-^* AngII -IAA), ranked by *p*-value. n = 5 mice/group. See **Fig. S3** for top enriched GO biological process and KEGG pathways for shared upregulated genes between genotypes (WT AngII-DTA and WT AngII-IAA; *Rag1^-/-^*AngII-DTA and *Rag1^-/-^*AngII-IAA), and between regions (WT AngII-DTA and *Rag1^-/-^* AngII-DTA; WT AngII-DTA and *Rag1^-/^* AngII-IAA).

The *Rag1^⁻/⁻^* aorta demonstrated amplified transcriptional response to AngII compared to WT. In *Rag1^⁻/⁻^*-DTA,17.55% of genes were differentially expressed, with upregulation of matrix remodeling genes (*Mmp2*, *Col1a1*), chemokines (*Ccl2*, *Ccl5*), and adipogenic regulators (*Pparg*, *Adipoq*), alongside suppression of *Nox4*, an oxidative stress enzyme integrating inflammatory cues with metabolic responses^23,24^. (**Fig. 3b**). *Rag1^⁻/⁻^*-IAA exhibited a 6.96% DEG response, marked by upregulated immune (*Lcp2*, *Cd44*) and metabolic (*Elovl6*, *Retn*) genes, among others (**Fig. 3c**). GO enrichment analysis in *Rag1^⁻/⁻^* mice revealed altered inflammatory signatures in the absence of adaptive immunity. Namely, in *Rag1^⁻/⁻^*-DTA, 8 of the top 15 upregulated pathways were immune-related (*negative regulation of immune system process,* etc.), suggesting potential innate immune compensation (purple; **Fig. 3d**). Conversely, the most significantly downregulated pathways of *Rag1^⁻/⁻^*-DTA centered on cardiovascular development and contractility (*aorta development*, *muscle contraction*, etc.), indicating suppression of structural and functional biological programs critical for aortic homeostasis (**Fig. 3d**). Similar to WT-IAA, the *Rag1^⁻/⁻^*-IAA appeared to preferentially express pathways vascular remodeling (*angiogenesis*, *extracellular matrix organization,* etc.) (**Fig. 3e**), aligning with its enhanced distensibility.

Hallmark pathway analysis confirmed distinct regional differences in gene expression patterns and biological responses between genotypes after AngII treatment. WT AngII DTA exhibited significant enrichment in both angiogenesis and inflammatory response pathways. In contrast, *Rag1^⁻/⁻^* AngII IAA demonstrated significant enrichment in adipogenesis and fatty acid metabolism pathways (**Fig. S2c**). GSEA plots further validated this finding, showing significant positive enrichment of adipogenesis pathways specifically in *Rag1^⁻/⁻^*AngII IAA samples (NES: 1.48, *p* < 0.001; **Fig. S2f**). WT AngII IAA and *Rag1^⁻/⁻^* AngII DTA samples showed no significant enrichment in these pathways (**Fig. S2d,e**). Together, these results suggest a unique, regionally-restricted metabolic reprogramming response in lymphocyte-deficient aortic tissue following AngII treatment.

Intersectional analysis of upregulated genes identified distinct transcriptional programs across genotypes and AngII treatment (**Fig. 3f**; **Fig. S3**). Notably, 1,977 genes (44.8% of DEGs across all groups) were uniquely upregulated in *Rag1*^⁻/⁻^AngII DTA – a 2.8-fold increase over WT AngII DTA (746 genes, 16.9% of DEGs) – highlighting the substantially elevated transcriptional activity of the thoracic aorta in the absence of adaptive immunity. The intersection of upregulated genes between *Rag1^⁻/⁻^* regions (*Rag1^⁻/⁻^*AngII DTA and *Rag1^⁻/⁻^*AngII IAA; 385 genes; 8.7% of DEGs) revealed significant enrichment in metabolic dysregulation, particularly in adipogenic and thermogenic pathways that were absent in WT mice. GO analysis of overlapping *Rag1^⁻/⁻^* DEGs identified enrichment in *temperature homeostasis* (GO:0001659, *p* = 4.90×10^-7^), *brown fat differentiation* (GO:0050873, *p* = 4.90×10^-7^), and *PPAR signaling* (KEGG:03320, *p* = 5.84×10^-7^) (**Fig. 3g**, **Fig. S3b**). Key drivers of this signature, including *Pparg*, *Adipoq*, *Fabp4*, and *Cd36*, are central to adipose metabolism and thermogenesis.

This transcriptomic analysis revealed distinct mechanisms of aortic remodeling in response to hypertension. While remodeling of the DTA appears to be dominated by T cell-driven inflammation, the IAA response is driven by structural adaptation mechanisms. Adaptive immunity restrained transcriptional responses to AngII, particularly in the DTA, while its absence in *Rag1*^⁻/⁻^ mice uncovered adipogenic and thermogenic biological programs across both lymphocyte-deficient aortic regions. These transcriptomic shifts implicate potential changes in PVAT remodeling and its functional role in modulating region-specific vascular dysfunctions during hypertension.

### Immune-Dependent PVAT Remodeling in Response to AngII Treatment

To define region-specific PVAT adaptations following AngII infusion, we analyzed lipid accumulation and PVAT area via Oil Red O staining in T-PVAT and A-PVAT depots (**Fig. 4a,d**). Quantification of positive red pixel area from brightfield microscopy images revealed distinct regional responses to AngII treatment. In WT mice, AngII significantly increased lipid content in T-PVAT (**Fig. 4b,c**; *p* < 0.05 vs. Saline), but not in A-PVAT (*p* > 0.05). In *Rag1*^⁻/⁻^ mice, PVAT area was slightly elevated over WT under saline conditions. Notably, *Rag1*^⁻/⁻^ T-PVAT lipid levels remained unchanged after AngII (*p* > 0.05), while A-PVAT showed a significant reduction in lipid content (**Fig. 4d,e**; *p* < 0.05 vs. Saline). Together, this suggests the absence of adaptive immunity reduced AngII-driven lipid expansion in T-PVAT, thus implicating lymphocyte-dependent inflammatory processes as a driver of pathological adiposity in WT T-PVAT. Conversely, A-PVAT atrophy in *Rag1*^⁻/⁻^ mice suggests either compensatory lipid catabolism or adipocyte apoptosis driven by a non-lymphocyte biological process.

**Fig. 4.**
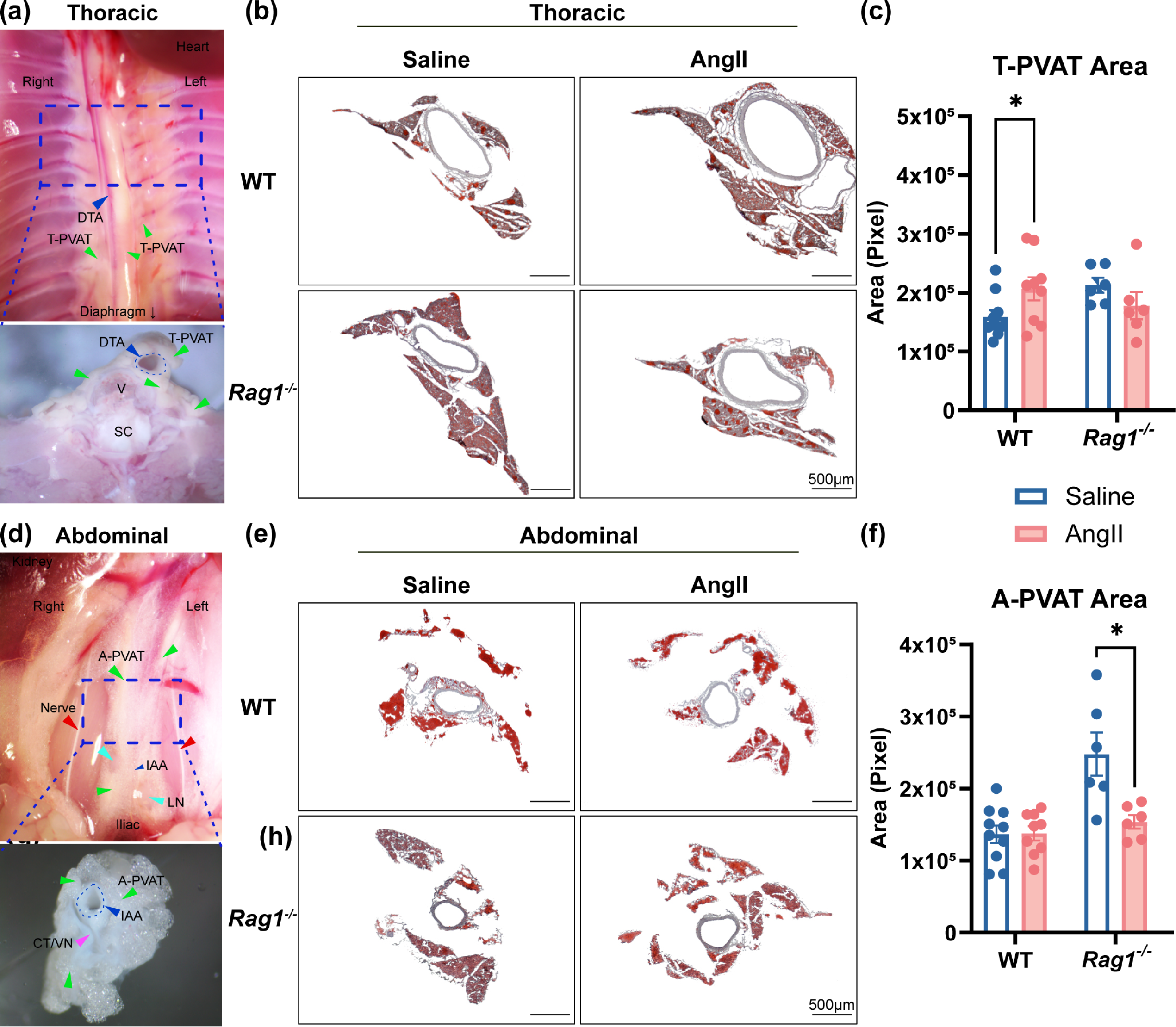
Lymphocyte-dependent lipid remodeling in perivascular adipose tissue. **(a)** Anatomic dissection images of thoracic perivascular adipose tissue (T-PVAT). Top-view and enlarged transverse section highlight PVAT (green arrowheads) adjacent to the DTA. Anatomical landmarks: V, vertebra; SC, spinal cord. **(b)** Representative Oil Red O staining of T-PVAT from WT and *Rag1^⁻/⁻^* mice following Saline or AngII treatment. **(c)** Quantification of lipid content in T-PVAT (positive red pixel area) following Saline (blue) or AngII (pink) treatment. **(d)** Anatomic dissection images of abdominal perivascular adipose tissue (A-PVAT). Top-view and enlarged transverse section highlight PVAT (green arrowheads) adjacent to the IAA. Anatomical landmarks: LN, lymph nodes; CT/VN, connective tissue/vein. **(e)** Representative Oil Red O staining of A-PVAT from WT and *Rag1^⁻/⁻^* mice following Saline or AngII treatment. **(f)** Quantification of lipid content in A-PVAT following Saline (blue) or AngII (pink) treatment. n = 10 mice for WT; n = 6 mice for *Rag1^⁻/⁻^*. Scale bar = 500µm. All data shown as mean ± SEM. Statistical significance within each genotype is from unpaired two-tailed Student’s *t*-tests for multiple comparisons denoted by \**p* ≤ 0.05.

### Immune-Metabolic Crosstalk Drives Regional PVAT Remodeling in Hypertension

To delineate mechanisms underlying PVAT remodeling during hypertension, we integrated profiling of ECM remodeling, inflammatory, and adipose-related genes with secretory proteomic analysis of PVAT-conditioned media to develop associations between region- and genotype-specific immune-metabolic crosstalk and region-specific AngII-induced vascular dysfunction. We first confirmed the identity of T-PVAT and A-PVAT depots using adipose phenotype-specific genetic markers. WT T-PVAT exhibited elevated expression of BAT-enriched genes, including *Ucp1* (*p* < 0.05 vs. A-PVAT, +0.93-fold in WT Saline and +0.85-fold in AngII groups; **Fig. 5a**) and *Pgc1a* (*p* < 0.05 in WT-AngII T-PVAT vs. A-PVAT; **Fig. S5a**). Notably, the difference in *Ucp1* expression between T-PVAT and A-PVAT was diminished in *Rag1^-/-^* mice (+0.24-fold *Rag1^-/-^* Saline T-PVAT vs. A-PVAT, +0.07-fold *Rag1^-/-^* AngII T-PVAT vs. A-PVAT), suggesting potential brown adipocyte alterations in *Rag1^-/-^* PVAT. In contrast, A-PVAT from *Rag1^⁻/⁻^* mice showed marked upregulation of WAT-specific genes, including *Adipoq* (*p* < 0.05 in *Rag1*^⁻/⁻^-Saline, *p* < 0.01 in *Rag1*^⁻/⁻^-AngII) and *Lep* (*p* < 0.001 in *Rag1*^⁻/⁻^-Saline, *p* < 0.05 in *Rag1*^⁻/⁻^-AngII), compared to the trending increase in *Ucp1* seen in T-PVAT (**Fig. 5a**). *Pparg* expression was significantly elevated in A-PVAT of *Rag1*^⁻/⁻^-AngII mice (*p* < 0.01; **Fig. S5a**). These findings indicate that T-PVAT retains BAT-like features in WT mice, while adaptive immune deficiency promotes a more pronounced WAT-like phenotype in A-PVAT. All markers of adipose phenotype, both in T-PVAT and A-PVAT, were largely unaffected by AngII treatment.

**Fig. 5.**
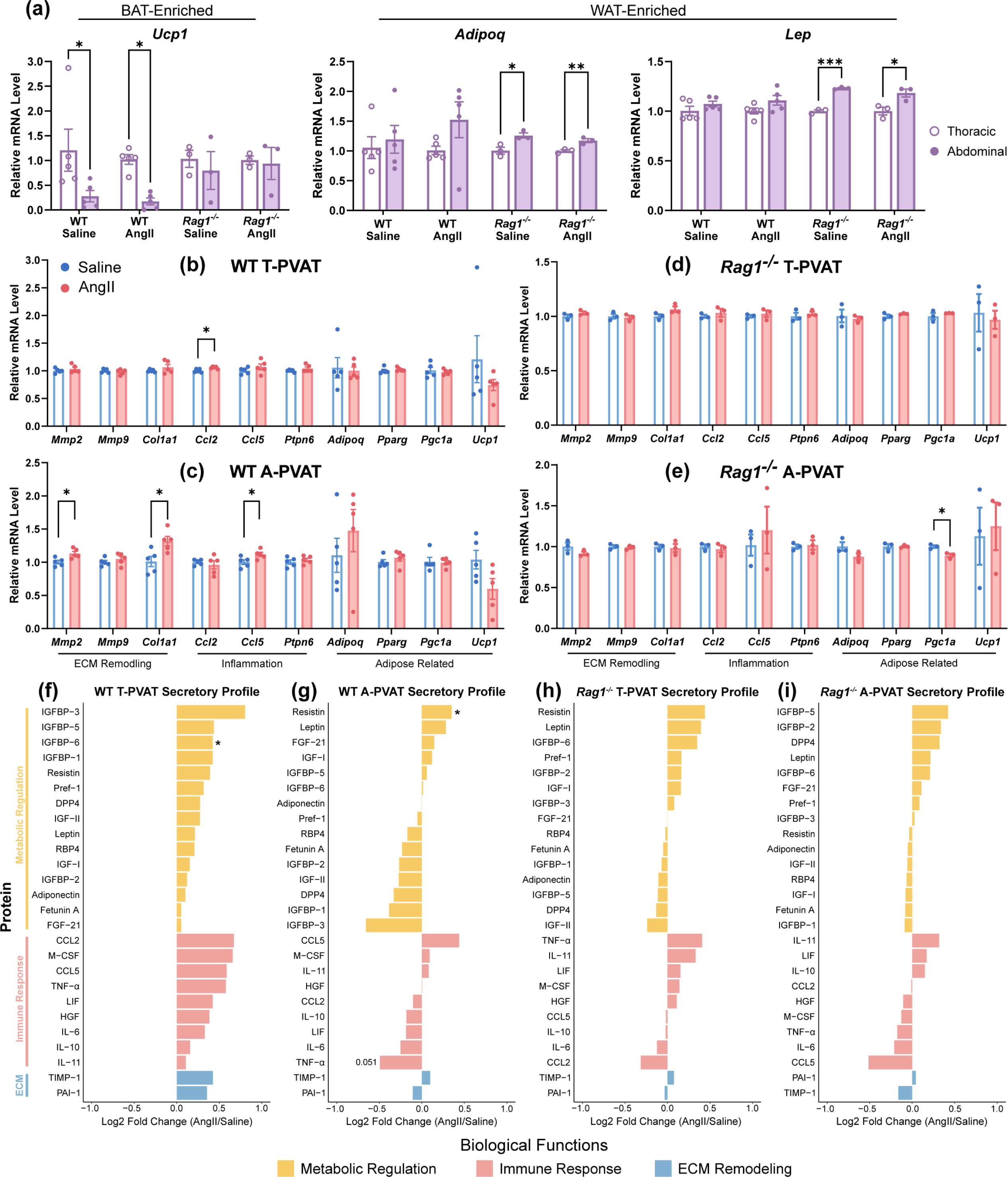
Immune-metabolic crosstalk drives region-specific PVAT remodeling. **(a)** Relative mRNA expression of adipose-specific genes canonically enriched in either brown adipocytes (*Ucp1*) or white adipocytes (*Lep, Adipoq*) from either T-PVAT (white) or A-PVAT (purple). **(b,c)** Transcriptional profiles of ECM remodeling, inflammatory and adipose-specific genes in **(b)** T-PVAT and **(c)** A-PVAT from WT mice following Saline (blue) or AngII (pink) treatment. **(d,e)** Corresponding gene expression in *Rag1^⁻/⁻^* **(d)** T-PVAT and **(e)** A-PVAT following Saline (blue) or AngII (pink) treatment. n = 5 mice for WT groups. n = 3 for *Rag1^⁻/⁻^* groups. **(f-i)** Adipokine array analysis of conditioned media from **(f,g)** WT PVAT and **(h,i)** *Rag1^⁻/⁻^* PVAT depots; **(f,h)** T-PVAT and **(g,i)** A-PVAT data shows relative expression following treatment by computing log2(fold change AngII/Saline). n = 3 mice/group. See **Fig. S5f** for representative membrane images and **Fig. S6** for all adipokine array results. All data shown as mean ± SEM. Statistical significance from unpaired two-tailed Student’s *t*-tests denoted by \**p* ≤ 0.05, \*\**p* ≤ 0.01, or \*\*\**p* ≤ 0.001.

Transcriptional profiling of ECM and immune-related genes revealed distinct remodeling patterns in response to AngII treatment. In WT mice, AngII induced significant upregulation of *Ccl2* in T-PVAT (*p* < 0.05 vs. Saline; **Fig. 5b**) and elevated *Mmp2*, *Col1a1*, and *Ccl5* in A-PVAT (*p* < 0.05; **Fig. 5c**). Fewer significant changes were observed in *Rag1*^⁻/⁻^ PVATs; only *Pgc1a* was altered after AngII (**Fig. 5d,e**; **Fig. S5d,e**). Notably, *Ucp1* expression trended downward in WT T-PVAT and A-PVAT after AngII, a pattern that was absent in *Rag1*^⁻/⁻^ mice, suggesting a potential T cell-dependent suppression of thermogenic biological processes in response to hypertension.

Adipokine array analysis of conditioned media revealed region- and genotype-specific secretory profiles (**Fig. 5f–i**; **Fig. S5f, S6**). In WT T-PVAT, there was a pronounced upregulation of protein secretion related to metabolic regulation (yellow; **Fig.5f**), immune response (pink; **Fig.5f**), and ECM (blue; **Fig.5f**). AngII enhanced expression of pro-inflammatory mediators (CCL5, M-CSF) and ECM regulators (TIMP-1), with IGFBP-6 showing a significant increase in WT T-PVAT (*p* < 0.05; **Fig. 5f**). Conversely, WT A-PVAT exhibited a divergent response. Resistin was significantly elevated (*p* < 0.05), meanwhile metabolic (adiponectin, DPP4) and immune-modulatory proteins (IL-10, TNF-α; *p* = 0.051) were suppressed (**Fig. 5g**). This paradoxical suppression of immune mediators in A-PVAT, despite its white adipose-like identity, highlights regional specialization in immunometabolic crosstalk that may contribute to regional differences in vascular remodeling. Lymphocyte deficiency profoundly reshaped PVAT secretomes. Notably, *Rag1^⁻/⁻^* mice exhibited blunted secretory responses with both regions showing attenuated inflammatory (CCL5) and metabolic (leptin) changes (*p* > 0.5; **Fig. 5h,i**). Moreover, the secretory profiles displayed trends toward discordant regulation (upregulated leptin vs. downregulated adiponectin) (**Fig. 5h,i**).

These findings demonstrate that AngII drives immune-dependent phenotypic shifts in PVAT, with T-PVAT adopting a pro-inflammatory, ECM-remodeling signature and A-PVAT exhibiting metabolic dysregulation. These observations are consistent with measured aortic biomechanical analyses (cf. **Fig.1**). Loss of adaptive immunity in *Rag1^⁻/⁻^* mice shifts T-PVAT away from a thermogenic profile toward a pro-inflammatory state and alters the expression of lipid-regulatory proteins in A-PVAT, suggesting that T cells help constrain depot-specific PVAT remodeling during hypertension.

### Immunofluorescence Analysis Reveals Region-Specific PVAT Phenotypic Plasticity

Immunofluorescence analysis of T-PVAT and A-PVAT established spatially divergent remodeling patterns in response to AngII treatment. In T-PVAT, AngII infusion significantly increased nuclei density in WT mice compared to Saline controls (1.2-fold, *p* < 0.05; **Fig. 6a,b**), a response that was lacking in *Rag1^⁻/⁻^* mice. The UCP1/Perilipin 1 (PLIN1) ratio – a compositional measure reflecting thermogenic dominance over lipid storage in PVAT – was reduced by 80% in *Rag1^⁻/⁻^*-Saline versus WT-Saline (*p* < 0.01; **Fig. 6c**). PLIN1-positive area, indicative of lipid-storing adipocytes, remained stable across genotypes and treatments in T-PVAT (**Fig. 6d**). UCP1 expression, marking thermogenic adipocytes, was suppressed by 67% in *Rag1^⁻/⁻^*-Saline compared to WT-Saline (*p* < 0.05; **Fig. 6e**). There was also a general, non-significant, trend toward increasing UCP1 expression in T-PVAT after AngII (+38.37% WT-AngII vs. WT-Saline, +141.91% *Rag1^⁻/⁻^*-AngII vs. *Rag1^⁻/⁻^*-Saline; **Fig. 6c,e**).

**Fig. 6.**
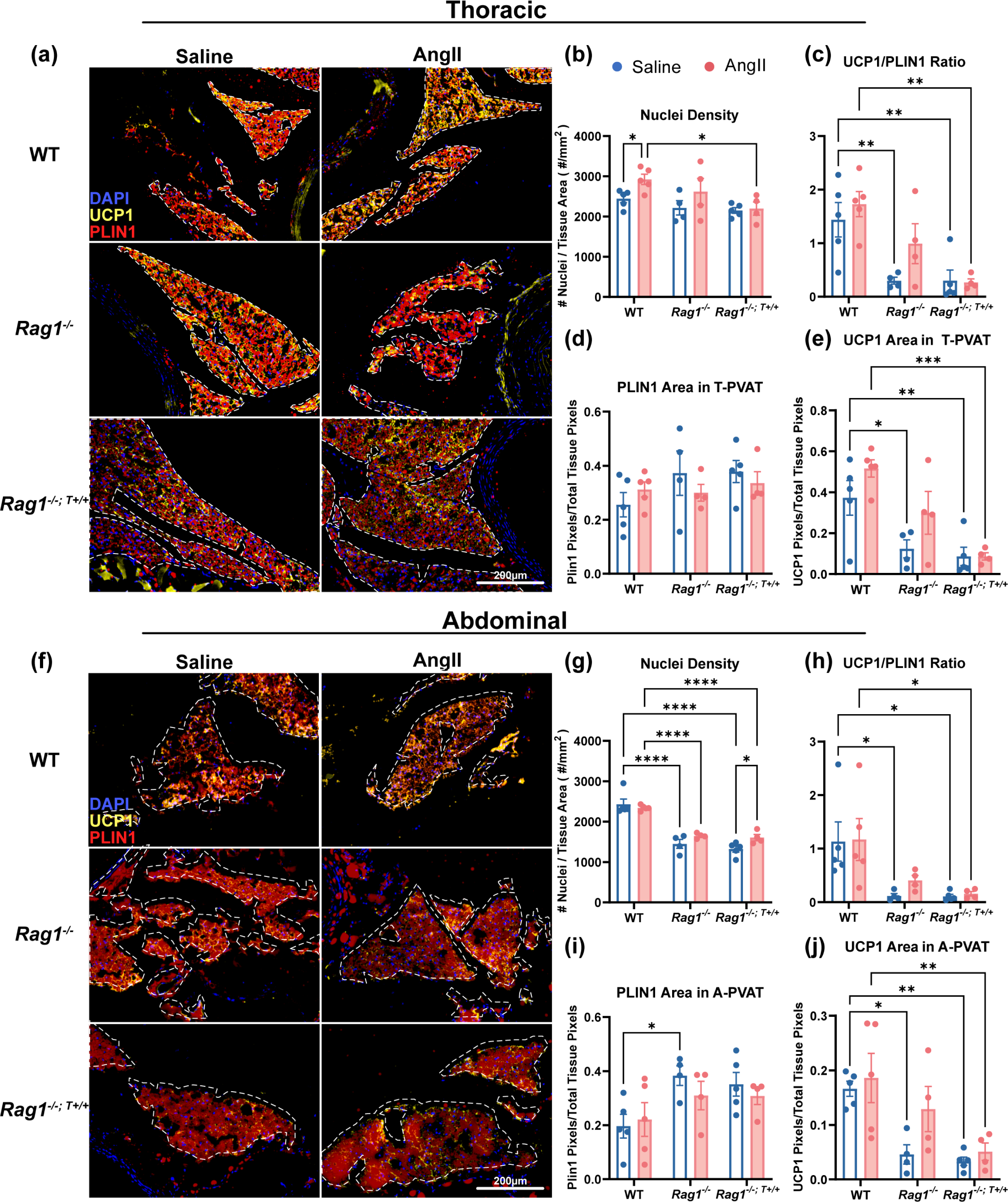
Adaptive immunity orchestrates region-specific PVAT phenotypic plasticity under hypertension. **(a)** Representative co-stained immunofluorescent images of UCP1 (yellow), perilipin-1 (PLIN1; red), and nuclei (DAPI; blue) in T-PVAT from WT, *Rag1^-/-^*, and *Rag1^-/-; T+/+^* (T cell reconstitution) mice. White dashed lines outline analyzed tissue regions. **(b–e)** Quantification of **(b)** nuclei density (number of cells per mm² of tissue), **(c)** UCP1/PLIN1 positive-stain-area ratio, **(d)** PLIN1-positive area (%), and **(e)** UCP1-positive area (%) in T-PVAT. **(f)** Representative A-PVAT immunofluorescence images. **(g–j)** Quantification of **(g)** nuclei density, **(h)** UCP1/PLIN1 positive-stain-area ratio, **(i)** PLIN1 area, and **(j)** UCP1 area in A-PVAT. n = 5 mice for WT. n = 4 mice for *Rag1^-/-^*. n = 5 mice for *Rag1^-/-; T+/+^*-Saline. n = 4 mice for *Rag1^-/-; T+/+^*-AngII. All data shown as mean ± SEM. Statistical significance from two-way ANOVA with post-hoc Tukey test for multiple comparisons denoted by \**p* ≤ 0.05, \*\**p* ≤ 0.01, \*\*\**p* ≤ 0.001, or \*\*\*\**p* ≤ 0.0001. See **Fig. S7** for all *p*-values for all effects in two-way ANOVA.

In A-PVAT, baseline nuclei density was markedly reduced in *Rag1^⁻/⁻^*-Saline versus WT-Saline (*p* < 0.0001; **Fig. 6g**), with no AngII-induced changes in either genotype. The UCP1/PLIN1 ratio was suppressed by 90% in *Rag1^⁻/⁻^*-Saline versus WT-Saline (*p* < 0.05; **Fig. 6h**). Meanwhile, *Rag1^⁻/⁻^*-AngII exhibited a non-significant 3.5-fold increase versus Saline. *Rag1^⁻/⁻^*-Saline mice exhibited a 95% elevation in PLIN1 area compared to WT-Saline (*p* < 0.05; **Fig. 6i**), suggesting expanded lipid storage in the absence of lymphocytes. UCP1 expression was suppressed by 72% in *Rag1^⁻/⁻^*-Saline versus WT-Saline (*p* < 0.05; **Fig. 6j**). Strikingly, *Rag1^⁻/⁻^*-AngII showed a 2.8-fold non-significant increase in UCP1 expression compared to *Rag1^⁻/⁻^*-Saline, indicating a trend toward thermogenic plasticity that may reflect compensatory adaptation in the absence of adaptive immunity.

### T Cell Reconstitution Restores Adaptive Immunity-Dependent PVAT Remodeling

Reconstitution of *Rag1^⁻/⁻^* mice with WT CD3+ T cells (*Rag1^⁻/⁻; T+/+^*) partially rescued region-specific phenotypes. In T-PVAT, T cell reconstitution attenuated AngII-induced cellular expansion, as shown by 75% lower nuclei density than WT-AngII (*p* < 0.05; **Fig. 6b**). While *Rag1^⁻/⁻^*-AngII exhibited a 3.3-fold increase in the UCP1/PLIN1 ratio compared to *Rag1^⁻/⁻^*-Saline (*p* < 0.05), reconstitution abolished this response (0.88-fold *Rag1^⁻/⁻; T+/+^*-AngII vs. *Rag1^⁻/⁻; T+/+^*-Saline; *p* > 0.05; **Fig. 6c**); however, UCP1 expression remained 77% lower in *Rag1^⁻/⁻; T+/+^*-AngII versus WT-AngII (*p* < 0.01; **Fig. 6e**), indicating incomplete thermogenic rescue. In A-PVAT, T cell reconstitution partially restored nuclei density and prevented AngII-induced UCP1 increases in *Rag1^-/-^* samples (*Rag1^⁻/⁻; T+/+^*-AngII vs. *Rag1^⁻/⁻; T+/+^*-Saline, *p* < 0.05; **Fig. 6g**). While *Rag1⁻^/⁻^*-AngII exhibited a 2.8-fold non-significant UCP1 elevation compared to Saline, reconstitution reduced this response to 1.5-fold in *Rag1^⁻/⁻; T+/+^*-AngII (**Fig. 6h,j**). Again, *Rag1^⁻/⁻; T+/+^*-AngII maintained significantly lower UCP1 expression than WT-AngII (*p* < 0.01; **Fig. 6j**), indicating adaptive immunity’s role in restraining AngII-induced thermogenesis.

### T Cell Reconstitution Partially Rescues Lipid Homeostasis and Vascular Remodeling

Oil Red O staining after T cell reconstitution revealed that adoptive transfer restored AngII-mediated lipid accumulation in T-PVAT with *Rag1^-/-; T+/+^* mice mirroring WT responses (cf. **Fig. 4c**) and exhibiting significant increases in lipid content compared to Saline controls (*p* < 0.05; **Fig. 7a,b**). On the other hand, A-PVAT lipid content in *Rag1^-/-; T+/+^* mice exhibited a non-significant reduction after AngII (**Fig. 7a,d**), which is consistent, yet attenuated, relative to the pronounced atrophy observed in *Rag1^⁻/⁻^*-AngII A-PVAT (cf. **Fig. 4f**).

**Fig. 7.**
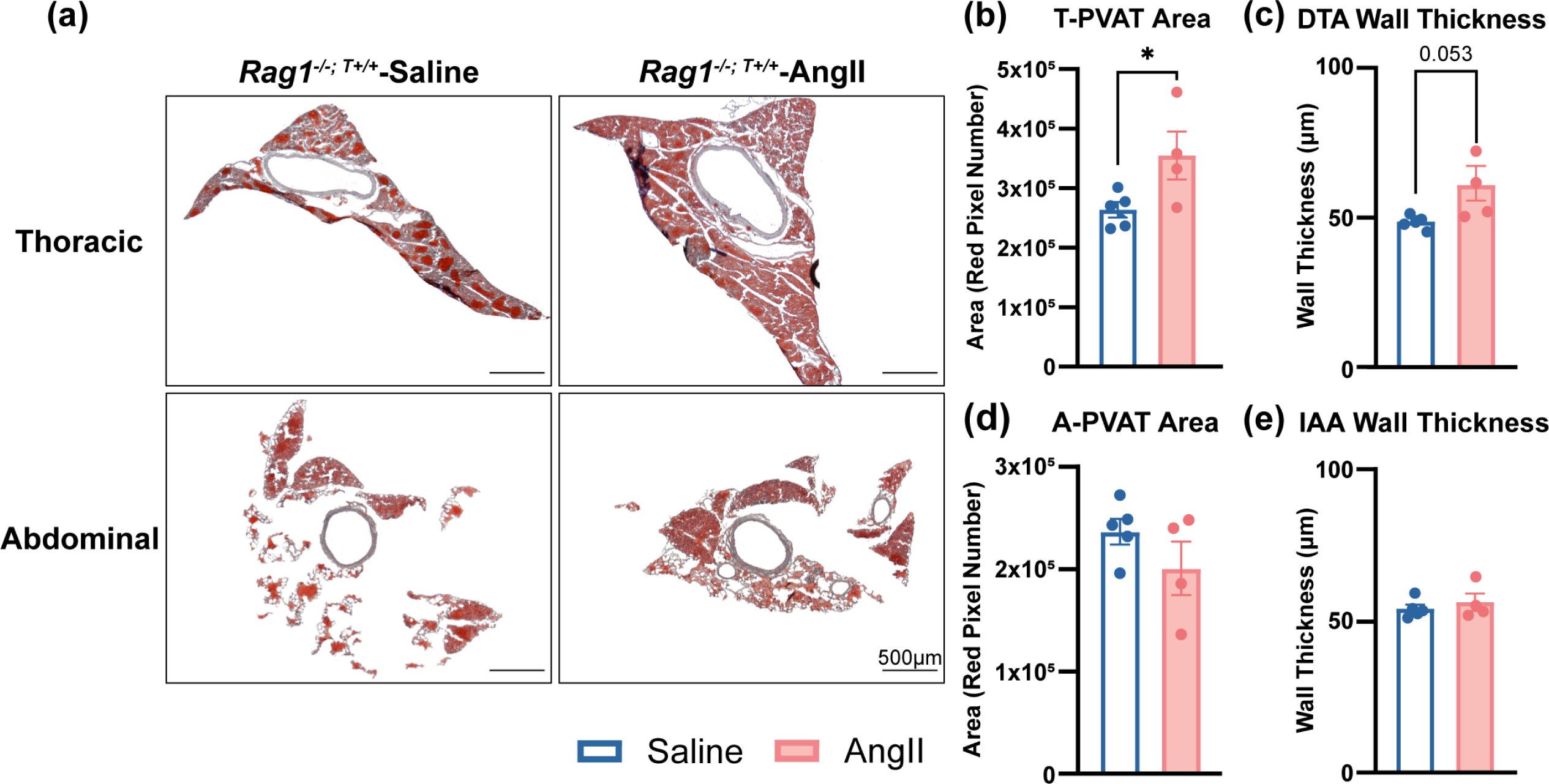
T cell reconstitution partially rescues lipid homeostasis and region-specific vascular remodeling in response to AngII. **(a)** Representative Oil Red O staining of T-PVAT and A-PVAT sections from *Rag1^⁻/⁻; T+/+^* mice treated with either Saline or AngII. **(b)** Quantification of lipid content in T-PVAT (positive red pixel area), **(c)** DTA wall thickness, **(d)** A-PVAT lipid content, and **(e)** IAA wall thickness. n = 5 mice for Saline. n = 4 for AngII. All data shown as mean ± SEM. Statistical significance from unpaired two-tailed Student’s *t*-tests denoted by \**p* ≤ 0.05.

T cell reconstitution also partially restored vascular remodeling patterns. *Rag1^-/-; T+/+^* mice displayed a strong trend toward increased wall thickness in the DTA after AngII (*p* = 0.053; **Fig. 7c**), nearly recapitulating the significant thickening seen in WT-AngII mice (*p* < 0.001; **Fig. S4a,c**). However, IAA wall thickness remained unchanged in *Rag1^-/-; T+/+^* mice after AngII administration (**Fig. 7e**), which was consistent with WT-AngII mice (minimal change) and divergent from *Rag1^⁻/⁻^*-AngII mice (significant thickening, *p* < 0.05; **Fig. S4b,d**).

These findings establish T cells as critical regulators of regional PVAT plasticity and subsequent regional vascular remodeling in response to AngII. Adaptive immunity promotes pathological lipid accumulation in T-PVAT while actively restraining thermogenic adaptation (i.e., browning) in A-PVAT. This dual regulation balances energy storage and expenditure across depots/vascular beds during hypertensive stimulus.

## Discussion

Hypertension drives heterogeneous vascular remodeling across aortic regions, but the mechanisms underlying this spatial disparity remain poorly understood. Our study establishes adaptive immunity as a pivotal regulator of region-specific aortic remodeling, with T cells amplifying maladaptation in the thoracic aorta while suppressing protective metabolic adaptions and browning in the abdominal aorta that enhance vascular compliance. By integrating biomechanical, transcriptomic, and histological analyses in WT and *Rag1^-/-^* mice, we demonstrate that PVAT heterogeneity and immune cell recruitment shape spatial disparities in vascular remodeling and dysfunction during hypertension. Together, these findings advance our understanding about the interplay between mechanical stress, immune activity, and metabolic adaptation in hypertensive vasculopathy, highlighting PVAT phenotype modulation as a potential target for manipulating aortic function.

AngII-induced hypertension triggered pronounced immune-mediated remodeling in WT-DTA, marked by wall thickening, reduced circumferential stiffness, and inflammatory gene upregulation (*Il6*, *Il17ra*, *Ccl2*). These changes align with prior reports of T cell-driven fibrosis in the thoracic aorta ^7,8^, but extend them by linking immune activity to PVAT’s role in regional susceptibility to fibrotic remodeling. The DTA is predisposed to T cell infiltration and ECM degradation due to its developmental origin of somite-derived mural cells with region-specific mechanosensitivity^7,25,26^. Infiltration of T cells in the perivascular space and adventitia has been associated with localized collagen accumulation and fragmentation of elastic laminae, both of which are key indicators of maladaptive vascular remodeling^6,7^. These findings mirror clinical observations in human thoracic aneurysms, where CD3^+^ T cell infiltration and IL-6 overexpression link to disease progression^27,28^. The concurrent suppression of the regulatory T cell (Treg) marker *Foxp3* observed in the current study suggests impaired resolution of inflammation may perpetuate thoracic aortic damage^29,30^. In *Rag1^-/-^* mice lacking mature lymphocytes, DTA remodeling was attenuated with near-normal wall thickness and preserved circumferential stiffness (cf. **Fig. 1a,f**). This underscores T cells’ necessity for maladaptation and positions adaptive immunity as a driver of thoracic aortic vulnerability, likely through compartmentalized immune-ECM crosstalk. Infiltrating T cells promote fibrosis via pro-inflammatory cytokines (e.g., IL-17), which induce collagen deposition^7,8^; suppression of Foxp3⁺ Tregs further impairs resolution of matrix disruption^29,30^. Tregs are key producers of transforming growth factor-beta (TGFβ), a master regulator of fibrillar collagen organization^31^. As a result, *Rag1^-/-^* mice exhibit attenuated remodeling with less changes in circumferential stiffness and wall architecture in thoracic region due to AngII treatment.

*Rag1^-/-^*-Saline mice exhibited reduced IAA distensibility (increased stiffness) versus WT, indicating that lifelong absence of adaptive immunity may alter baseline ECM composition. Strikingly, AngII infusion paradoxically reversed this phenotype in *Rag1^-/-^*-IAA by inducing significantly increased distensibility (decreased stiffness) relative to Saline controls, thus unmasking compensatory remodeling in the absence of lymphocytes. Critically, these compliance changes occurred exclusively in the circumferential direction, mirroring the direction-specific vulnerability in aortic aneurysms where inflammation-driven ECM degradation compromises circumferential integrity^6,9,25^.

While the IAA demonstrated resilience in WT mice, *Rag1^-/-^* mice exhibited altered distensibility and PPARγ-associated adipogenic reprogramming. Despite maintaining near-homeostatic wall thickness after AngII, *Rag1^-/-^*-IAA showed reduced circumferential stresses, suggesting potential smooth muscle-mediated buffering against mechanical perturbations^5,32,33^. In *Rag1^-/-^* mice, the absence of adaptive immunity unmasked latent metabolic pathways, including upregulation of *Pparg* and *Adipoq*, which are genes associated with improved vascular compliance^14^. PPARγ agonists reduce arterial stiffness by suppressing inflammation and enhancing nitric oxide bioavailability, suggesting that T cells may simultaneously constrain protective metabolic reprogramming (browning) in the abdominal aorta despite its inherent resilience^34^. These adipogenic shifts likely contribute to ECM remodeling, as supported by studies linking PPARγ/adiponectin to reduced collagen deposition in distal vasculature^35–37^. This metabolic alteration in distal aortic regions could thus mitigate hypertensive damage, positioning adaptive immunity as a critical determinant of regional susceptibility^5^.

The striking emergence of adipogenic and thermogenic gene signatures in *Rag1^-/-^* mice across both aortic regions further supports the idea of lymphocyte-mediated metabolic suppression. This metabolic reprogramming, driven in part by genes including *Pparg*, *Adipoq*, and *Fabp4*, was absent in WT mice, suggesting that adaptive immunity contributes to suppression of these pathways during hypertension. Transcriptomic analysis further revealed that the DTA exhibited a robust inflammatory signature while the IAA demonstrated a more restrained transcriptional response focused on ECM organization. These distinct transcriptional programs reflect divergent strategies for adaptation in response to excess mechanical loading during hypertension: inflammatory-driven structural remodeling in the thoracic region versus matrix-oriented turnover and reinforcement in the abdominal segment.

PVAT heterogeneity emerged as a pivotal regulator of region-specific vascular remodeling. In WT mice, AngII promoted lipid accumulation (increased area) and a pro-inflammatory secretory profile in T-PVAT characterized by significant upregulation of IGFBP-6 alongside elevated pro-inflammatory mediators (CCL5, M-CSF) and ECM regulators (TIMP-1). IGFBP-6 likely exacerbates thoracic aortic fibrosis and ECM degradation by sequestering other IGFs and enhancing TGF-β signaling ^31^, which could be a key factor contributing to the maladaptive biomechanical remodeling observed in the DTA. Concurrently, TIMP-1 overexpression promotes collagen stabilization and adventitial fibrosis, further compromising vascular compliance^5,33^. Paradoxically, despite T-PVAT’s brown adipose-like identity, this inflammatory skewing (e.g., CCL5-driven T cell recruitment^12^) appears to override its inherent thermogenic properties, creating a perivascular milieu that amplifies thoracic vascular dysfunction^12^. T-PVAT’s brown adipose tissue-like phenotype under basal conditions was skewed toward a pro-inflammatory state during hypertension despite slight increases in UCP1 protein expression (cf. **Fig. 6c,e**), thus exacerbating thoracic aortic damage through aberrant paracrine signaling. On the other hand, WT A-PVAT exhibited divergent immunometabolic reprogramming in response to AngII treatment. Resistin, a key driver of endothelial dysfunction and oxidative stress^10,20^, was significantly upregulated in WT-AngII A-PVAT, while immune-modulatory cytokines trended downward. This suggests A-PVAT, which is classically considered to be white-like, undergoes metabolic dysregulation rather than excess inflammation due, at least in part, to PPARγ suppression^13,19^. The observed resistin elevation (cf. **Fig. 5g**) aligns with A-PVAT’s reduced thermogenic plasticity and may contribute to impaired vasodilation, though the biomechanical resilience of the IAA persists due to compensatory ECM reorganization.

The observation that T-PVAT exhibits canonical brown adipose markers (e.g., *Ucp1*) yet drives pro-inflammatory maladaptive remodeling is inconsistent with the classical view of brown adipose as metabolically protective and anti-inflammatory^13,16,17,38^. While thermogenic adipose depots typically mitigate metabolic dysfunction via anti-inflammatory cytokines (e.g., IL-10) and vasodilatory adipokines (e.g., adiponectin)^10,15^, T-PVAT’s perivascular niche and immunological crosstalk may override these beneficial characteristics under hypertensive stimulus. Similar contradictions can be found in other contexts: for example, *Ucp1^+^* adipocytes present in the crown-like structures of obese visceral fat exhibit excess inflammation despite maintaining their thermogenic identity^22^, and PVAT in various models of atherosclerosis has been shown to exhibit concurrent thermogenic activation (browning) and oxidative stress^21^. This suggests anatomical location, immune microenvironment, and disease-specific cues could possibly decouple thermogenic capacity from the anti-inflammatory functions of brown adipocytes and thereby alter PVAT function.

On the other hand, *Rag1^-/-^* mice exhibited a non-statistically significant tendency toward thermogenic plasticity in A-PVAT, characterized by modest *Ucp1* upregulation and reduced lipid content (cf. **Fig. 5a,e** and **Fig. 6f,i**). This compensatory thermogenic adaptation, unmasked in the absence of adaptive immunity and attenuated by T cell reconstitution (cf. **Fig.7**), suggests adaptive immunity suppresses PVAT’s capacity for metabolic adaptation^39^ potentially preventing excessive phenotypic shifting during transient stressors. The anatomical gradient in PVAT phenotype – from brown-like in the thoracic region to white-like in the abdominal region – provides a molecular basis for the observed regional differences in immune cell recruitment and vascular remodeling. Thus, adaptive immunity acts as a key regulator not only for ECM homeostasis but also for PVAT phenotypic plasticity. This dual regulation may reflect an evolutionary trade-off. By tempering metabolic overcompensation, T cells could stabilize vascular function during acute insults, though at the cost of reducing long-term adaptive reserves in chronic disease.

Adipose browning has been associated with improved vascular function through vasodilatory metabolite secretion and enhanced energy expenditure^13^, aligning with studies showing that PVAT-derived factors mitigate endothelial dysfunction in hypertension^15,38^. The suppression of thermogenic markers (UCP1) in *Rag1^-/-^* mice under basal conditions, coupled with their paradoxical increase under AngII treatment, reveals a complex interplay between adaptive immunity and adipose tissue phenotype and energy expenditure. These findings suggest adipose browning, particularly in A-PVAT in response to hypertensive stimulus, may represent a compensatory regenerative mechanism. Recent research has recognized adipose browning for its regenerative potential beyond simple metabolic enhancement^38,40,41^. Thermogenic activation (browning) of adipose promotes paracrine tissue repair through enhanced angiogenesis and anti-inflammatory cytokine production^42,43^. The suppression of this regenerative browning by adaptive immunity thus represents a previously unrecognized mechanism by which the immune system may exacerbate vascular dysfunction during hypertension.

While our findings provide critical insights into the role of adaptive immunity and PVAT heterogeneity in region-specific aortic remodeling, several limitations need to be acknowledged and taken into consideration. First, sex-specific differences in hypertensive remodeling were not addressed, as we focused on male mice to isolate T cell-mediated mechanisms. Estrogen’s vasoprotective effects, including suppression of vascular inflammation and oxidative stress, likely mask, or at least mitigate, immune-driven remodeling responses in females^44,45^. Future studies comparing sexes could clarify whether PVAT’s thermogenic plasticity is estrogen-dependent, thus offering insights into sex disparities in hypertensive complications. Additionally, the lifelong absence of mature T and B cells from *Rag1^-/-^* mice potentially introduces developmental adaptations that can influence vascular remodeling and PVAT responses. T cells, particularly regulatory subsets such as Tregs, play established roles in adipose tissue development, homeostasis, and browning capacity^39^. Thus, the impact of *Rag1* deficiency on Tregs, key promoters of beige adipogenesis via IL-33/IL-4 signaling^39,46^, could fundamentally alter PVAT phenotype and baseline metabolic potential thereby confounding interpretations of acute hypertensive responses. Inducible cell-type-specific knockout models could mitigate this developmental confounder. Third, although our AngII model recapitulates acute hypertensive remodeling, chronic studies are needed to assess long-term outcomes. Prolonged PPARγ activation in *Rag1^-/-^* mice may paradoxically drive fibrosis, as observed in metabolic disorders, necessitating further functional analyses of PVAT-derived metabolites on vascular cells^47,48^. Furthermore, our adoptive transfer approach utilized total CD3^+^ T cells which precludes the delineation of contributions from specific T cell subsets (e.g., CD4^+^ vs. CD8^+^, Th1/Th17 vs. Tregs) to regional PVAT remodeling and vascular dysfunction. Given the opposing roles of pro-inflammatory (e.g., Th17) and regulatory T cells in hypertension^7,8,20,29,30^, future studies employing subset-specific transfers or depletions are essential.

The incomplete partial rescue of phenotypes by CD3^+^ T cell reconstitution indicates potential contributions from other lymphocyte populations absent in *Rag1^-/-^* mice, such as B cells or innate lymphocytes^49,50^. For example, B cell-derived IL-10 is known to modulate vascular inflammation in hypertension^50^, but how it interplays with T cells in regional remodeling remains unexplored. Computational fluid dynamics could further elucidate how altered shear stress in thoracic versus abdominal aortas influences T cell recruitment and PVAT phenotype^51–53^.

In conclusion, our work contributes to the redefinition of hypertension and hypertensive vascular remodeling as a disorder of immune-mechano-metabolic integration^54^, modulated by PVAT heterogeneity. Adaptive immunity promotes maladaptive thoracic remodeling through T cell-driven inflammation and ECM degradation, while restraining compensatory thermogenic reprogramming (browning) in the abdominal aorta. PVAT serves as a critical interface wherein immune activity modulates adipose plasticity to influence vascular tone and ECM deposition in a region-specific manner. The distinct molecular and functional characteristics of T-PVAT versus A-PVAT create microenvironments that differentially influence local vascular remodeling in response to hypertension. Targeting T cell-PVAT crosstalk or enhancing thermogenic plasticity may preserve aortic compliance without exacerbating inflammation, offering spatially resolved strategies to mitigate hypertensive vascular remodeling and disease.

## Acknowledgements

We thank the Genome Technology access Center at the McDonnell Genome Institute at Washington University School of Medicine for assistance with genomic analysis. The Center is partially supported by NCI Cancer Center Support Grant #P30 CA91842 to the Siteman Cancer Center and by ICTS/CTSA Grant# UL1TR002345 from the National Center for Research Resources (NCRR), a component of the National Institutes of Health (NIH), and NIH Roadmap for Medical Research. This publication is solely the responsibility of the authors and does not necessarily represent the official view of NCRR or NIH. Additionally, we would also like to acknowledge the lab of Srikanth Singamaneni, PhD in the Department of Mechanical Engineering and Materials Science at Washington University in St. Louis for providing access to their plasmonic fluor technology and support with adipokine membrane imaging. Finally, this work was supported, in-part, by grants from the National Institutes of Health (R00 HL146951 to MRB), the American Heart Association (941138 to MRB), and the National Science Foundation (2340666 to MRB).

## Author Contributions

Y.X. and C.S. designed the research. Y.X. performed experiments, analyzed data, and wrote the manuscript. M.R.B. conceived of experimental designs and supervised the project. All authors reviewed and approved the final version of the submitted manuscript.

## Data Availability

The RNA-seq data have been deposited in the Gene Expression Omnibus database (GEOXXX).

